# Accuracy, limitations and cost-efficiency of eDNA-based community survey in tropical frogs

**DOI:** 10.1101/176065

**Authors:** Miklós Bálint, Carsten Nowak, Orsolya Márton, Steffen U. Pauls, Claudia Wittwer, B. Jose Luis Aramayo, Arne Schulze, Thierry Chambert, Berardino Cocchiararo, Martin Jansen

## Abstract

Rapid environmental change in highly biodiverse tropical regions demands efficient biomonitoring programs. While existing metrics of species diversity and community composition rely on encounter-based survey data, eDNA recently emerged as alternative approach. Costs and ecological value of eDNA-based methods have rarely been evaluated in tropical regions, where high species richness is accompanied by high functional diversity (e.g. the use of different microhabitats by different species and life-stages). We first tested whether estimation of tropical frogs’ community structure derived from eDNA data is compatible with expert field assessments. Next we evaluated whether eDNA is a financially viable solution for biodiversity monitoring in tropical regions. We applied eDNA metabarcoding to investigate frog species occurrence in five ponds in the Chiquitano dry forest region in Bolivia and compared our data with a simultaneous visual and audio encounter survey (VAES). We found that taxon lists and community structure generated with eDNA and VAES correspond closely, and most deviations are attributable to different species’ life histories. Cost efficiency of eDNA surveys was mostly influenced by the richness of local fauna and the number of surveyed sites: VAES may be less costly in low-diversity regions, but eDNA quickly becomes more cost-efficient in high-diversity regions with many sites sampled. The results highlight that eDNA is suitable for large-scale biodiversity surveys in high-diversity areas if life history is considered, and certain precautions in sampling, genetic analyses and data interpretation are taken. We anticipate that spatially extensive, standardized eDNA biodiversity surveys will quickly emerge in the future.

## Introduction

Improvements on most biodiversity loss indicators lag behind the 20 “Aichi Biodiversity Targets” (UNEP 2016) that aim to reduce the decline of biodiversity by 2020 (Tittensor *et al*. 2014). An important component of the biodiversity crisis is the extinction of species. Based on current trends in mammals, birds, reptiles and amphibians, it has been projected that the biodiversity crisis may lead to the 6th Mass Extinction over the next three centuries if all threatened species go extinct (Barnosky *et al*. 2011). Current rates of extinction may even be much higher if one considers the extinction that likely occurred during the last few decades-centuries, but went unnoticed because the now-extinct species had small ranges, were never described or only described on the eve of their extinction (Pimm *et al*. 2014; Lees & Pimm 2015). It is however difficult to assess which species are endangered and to what extent. First, most taxa remain poorly described: in some highly diverse regions many species will likely go extinct before they are discovered (Costello *et al*. 2013; Lees & Pimm 2015). Second, cryptic genetic diversity is common within morphospecies (Pfenninger & Schwenk 2007; Pauls *et al*. 2013), and global change may impact cryptic diversity more severely than morphospecies (Bálint *et al*. 2011). Third, data on population-level trends is scarce, even for well-known species (Butchart *et al*. 2010). Better population-level biodiversity data is thus urgently needed to 1) understand biodiversity patterns and extinction threats, 2) improve forecasting abilities about future biodiversity, and 3) improve humanity’s responses to the challenges of biodiversity loss. This data is crucial in times when conservation action is increasingly demanded by society (Tittensor *et al*. 2014).

The importance of internationally coordinated, standardized biodiversity data collection is long recognized both in science and in conservation (Henry *et al*. 2008). This is particularly true for the most biodiverse areas. The tropics are generally underrepresented in ecological studies (Clarke *et al*. 2017; Stroud & Feeley 2017). In addition, encounter-based data collection is logistically challenging since it needs considerable funds to bring enough specialists for different organismic groups to remote areas, thus insufficient funds and expertise often limit comprehensive surveys. Indirect species records through environmental DNA are increasingly heralded as an alternative to encounter-based surveys (Thomsen & Willerslev 2015; Pedersen *et al*. 2015). eDNA also facilitates the standardization of biodiversity surveys at both regional and global scales since community composition data can be obtained by high-throughput-sequencing of standardized, taxonomically informative marker genes (metabarcoding) (Taberlet *et al*. 2012a; Cristescu 2014). Aquatic or semiaquatic vertebrates such as frogs and other amphibians or fish have been early targets of eDNA based studies, either focusing on the detection of single species (e.g. Ficetola *et al*. 2008; Goldberg et *al*. 2011; Jerde *et al*. 2011; Thomsen *et al*. 2012a) or entire communities (e.g. Thomsen *et al*. 2012b; Valentini *et al*. 2016; Shaw *et al*. 2016; Yamamoto *et al*. 2017). The use of next-generation sequencing approaches led to a boost in data acquisition (Taberlet *et al*. 2012b) and is considered to make important contributions to biodiversity research (Rees *et al*. 2014; Bohmann *et al*. 2014). eDNA-based metabarcoding may present one of several tools needed to globally coordinate initiatives for ecosystem monitoring and sustainable management (Bush *et al*. 2017; Schmeller *et al*. 2017).

In this study we evaluate whether eDNA metabarcoding is suitable for inventories of frogs, a group with particular high species-diversity in tropical regions. Frogs and other amphibians are sentinel victims of the biodiversity crisis: more than one-third of the approximately 7500 described species are endangered (Stuart *et al*. 2004; Bishop *et al*. 2012; Whittaker *et al*. 2013). Frogs are also known for being a highly diverse, but incompletely described taxon, especially in the tropics (Ferrão *et al*. 2016; Caminer *et al*. 2017). Many “widespread” morphospecies harbor considerable cryptic genetic diversity and are better considered complexes of closely related species with much smaller ranges (Fouquet *et al*. 2007; Gehara *et al*. 2013, 2014; Ortega-Andrade *et al*. 2015). Efficient implementation of amphibian conservation measures (e.g. the prioritization of areas for conservation, or informing society and stakeholders about conservation needs) are only possible with geographically broad-scaled fine-grain, taxonomically well-resolved faunistic data, but our current understanding of present and future amphibian biodiversity is often based on rare, spatially and temporally scattered observations of phenotypically defined taxa.

First efforts have been made to test the suitability of eDNA for the survey of tropical frog biodiversity (Lopes *et al*. 2016), but important practical aspects remain unaddressed. First, it is not clear which fraction of the local species pool is represented by amphibian eDNA in tropical water bodies. Existing comparisons of encounter-based surveys and eDNA either do not include non-adult life stages, or they use already compiled fauna lists for the evaluation of eDNA performance without consideration of life history traits or behavioral aspects at the moment of sampling. However, temporal and spatial patterns of microhabitat use by frogs is largely species-specific and influenced by phenology and reproduction modes (e.g. Duellmann & Pyles 1983, Haddad & Prado 2005, Wells 2010), which can induce strong biases in biodiversity data (Petitot *et al*. 2014). Second, the degree to which aquatic eDNA can provide accurate assessments of community structure remains largely unevaluated. Most studies to date have only investigated the correspondence between encounter-based and metabarcoded taxon lists (Miya *et al*. 2015; Valentini *et al*. 2016), although community structure assessments themselves may be of higher importance for many applications (Ji *et al*. 2013; Elbrecht *et al*. 2017). Third, it is not clear whether, and under what conditions eDNA is financially efficient since comparisons are lacking (Lopes *et al*. 2016), although these comparisons are essential for deciding on data collection strategies.

Here we address whether 1) detectability of tropical amphibians with eDNA is linked to species’ life history, and 2) eDNA sampling provides accurate data for the characterization of community assembly. Finally, we present a framework for cost comparisons between encounter- and eDNA-based biodiversity survey that may be adapted to other systems beyond amphibians. We compare the results of long- and short-term encounter-based field surveys, and an eDNA survey of tropical amphibians in a well-characterized high-diversity area in Bolivia.

## Materials and methods

The study area is located the Chiquitano region of Bolivia, a forest-savanna ecotone between Amazon, Cerrado and Gran Chaco in a transition zone among humid and dry forests that are special in regard to their taxonomic and functional diversity (Castro *et al*. 1999). The region contains the largest intact, old-growth block of seasonally dry tropical forests in South America (Miles *et al*. 2006; Power *et al*. 2016). Samples were collected from ponds in the vicinity of the Biological Station “Centro de Investigaciones Ecológicas Chiquitos” on the San Sebastián cattle ranch (S16.3622, W62.00225, 500 m a.s.l.), 24 km south of the town of Concepción, Province of Ñuflo de Chávez, Santa Cruz Department, Bolivia. A description of the area is given by Schulze *et al*. (2009). This area is well characterized with respect to amphibians, including both larvae and adults (Schulze *et al*. 2015). Intensive long-term assessments have resulted in the detection of 45 frog species in the area (e.g. Jansen 2009; Schulze *et al*. 2009, 2015; Jansen *et al*. 2011), as well as the discovery of cryptic intraspecific diversity (e.g. Jansen *et al*. 2011; Jansen *et al*. 2016).

We sampled five ponds (T1 – T5) for this study between 19 and 23 November 2014 (see also section “Site and ponds” in the Supplemental information). At these ponds, 35 species have been recorded in previous long-term surveys. Water samples were obtained from three sampling points at each pond (Fig. 1): These water samples consisted of four liters of water collected into two two-liter silicon bottles. One sample of 2 × 2 liters was taken at each sampling point of ponds T1 - T4. Three samples of 2 × 2 liters were taken on each sampling points of pond T5 to check for sampling variation, i.e. whether replication in the sampling records more variation in community composition compared to replication in PCR steps. We also sampled and filtered water from two field controls: Tap water from the station from a covered well, and water from an aquarium with tadpoles of *Leptodactylus vastus* at the station. The water samples were filtered at the station immediately after collection.

**Fig. 1.**
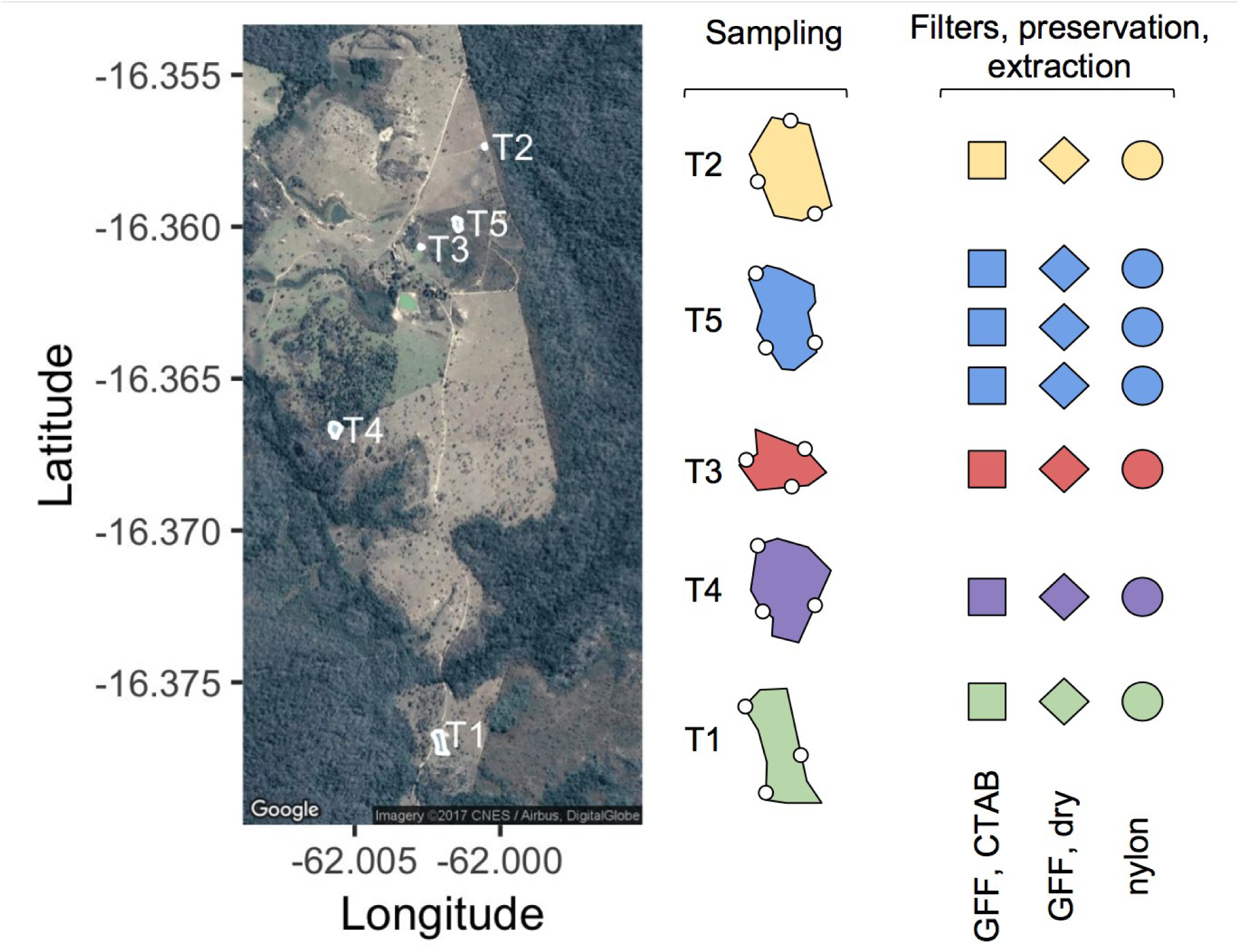
Sampled ponds, sampling replication, and filtration - filter preservation strategy. GFF - glass-fiber filter, 2 μm; nylon - nylon filter, 0.2 μm. Each sample (three per pond, marked with small circles) were processed with each three filtration strategies. Multiple samples of pond T5 were three times processed with each filtration strategy.

Samples were filtrated with a vacuum filtration unit (Duran Group, Wertheim, Germany), which was connected to a vacuum membrane pump (Laboport, both Carl Roth GmbH + Co. KG, Karlsruhe, Germany). Filtration was performed until complete clogging of the filters. For each sample, the water from the two bottles was filtered separately on two glass-fiber filters (GFF, pore size 2 μm). One of the filters was preserved in CTAB (A), the other filter was dried immediately (B). Our intention here was to evaluate if filter preservation influences species detection, since the transportation of dry filters may be considerably simpler compared to wet filters. The flow-through from the two GFF filtrations was combined and filtered with a nylon filter until clogging (NF, pore size 0.2 μm, Millipore, Merck, Darmstadt, Germany). This filter was immediately dried after filtration (C). We intended to evaluate if the larger pore-size filters sufficiently capture anuran eDNA, or finer pore filters allow for additional species detection. To avoid cross-contamination we changed gloves with every new sample and disinfected hands with ethanol (96%), as well we cleaned filtration unit with ethanol (96%) after each pond or site.

Combined visual and audio encounter surveys (VAES, Zimmerman 1994), and a tadpole survey (TS, Schulze *et al*. 2015) were conducted by experienced observers (MJ and AS) to generate a presence/absence matrix of species for each pond. Frogs surveys were performed at night (between 21:00 and 00:30 hours) during 0.5 - 1 hrs transect walks along the ponds to detect and identify specimens in vegetation, in water and on the ground around ponds. Detections occurred from either visual (using flashlights and headlamps) or auditory species identification. Tadpoles were sampled with dip nets once during daytime and once at night in each pond as well as the riparian vegetation.

eDNA samples were processed in Germany in a laboratory dedicated to the pre-PCR handling of environmental DNA samples. Several working routines have been implemented in the laboratory to avoid contamination of samples and reactions, including spatial separation from the other DNA facilities (separate room), strict decontamination protocols using UV light and bleach, physical separation of extraction and PCR work spaces, automated extraction, and restricted access to staff trained in the handling and analysis of forensic and environmental samples.

DNA was extracted from GFF samples (A, B, see above) with a CTAB chloroform extraction method according to Strand *et al*. (2014) and Wittwer *et al*. (2017). Dried nylon filters (C) were extracted with DNeasy Blood & Tissue Kit (Qiagen, Hilden, Germany) following Thomsen *et al*. (2012b). Negative controls were included in all extraction sessions (see also DNA extraction in the Supplemental information). The barcode amplification targeted mitochondrial 16S ribosomal DNA. We designed primers (see Primer design in Supplemental information for details on design and primer tests) for a 150 bp fragment with the reference database of local species (sequences from 159 specimens, Jansen *et al*. 2011; Schulze *et al*. 2015). The primers were tested on a subset of 12 species. PCRs were run on 96-well plates in 15-μl reaction volumes with a touchdown protocol, with four technical PCR replicates per sample (see PCR setup in the Supplemental information for PCR protocol and cycling conditions). We included four negative PCR controls (ultra-sterile water), four extraction blanks, and two positive controls on each plate. Positive controls consisted of 1) equimolar positives where DNA of the twelve frog species (primer development) was pooled in equimolar concentration (5 ng/μl), and 2) non-equimolar positives where a concentration series was used by diluting the 5 ng/μl templates 1x to 512x dilution. PCR replicates were individually labeled with multiplexing indices designed by Kozarewa and Turner (2011). PCR products were purified with Agencourt AMPure XP SPRI magnetic beads (Beckman Coulter, Brea, CA), pooled in equal volumes, and paired-end-sequenced (2 × 150 bp) on an Illumina NextSeq 500 sequencer at StarSEQ GmbH (Mainz, Germany).

The bioinformatic pipeline for sequence processing mostly relied on OBITools v1.2.0 (Boyer *et al*. 2016, see Sequence processing in the Supplemental information for details). We performed two reference-based taxonomic assignments, first with a custom database of 16S sequences of 159 specimens from the regional amphibian fauna (16S_custom), and second with all 16S sequences found in the EMBL (release 125, 16S_EMBL). Further filtering was done in R (R Core Team 2017) with a script supplied on GitHub: https://github.com/MikiBalint/amphibian-eDNA. This includes an ordination of all PCR replicates, negative and positive controls. This ordination was used to identify outlier replicates of samples from different ponds (Supplemental Fig. S1).

We followed recommendations for the statistical analyses of marker gene community data (Bálint *et al*. 2016). eDNA faunistic differences among the five ponds were visualized with a latent variable model-based ordination in R (‘boral’, Hui *et al*. 2015), and tested with multispecies generalized linear models (GLMs in ‘mvabund’, Wang *et al*. 2012). We used read abundances as input data for the boral ordination and the multispecies GLMs since presence-absences inferred from read abundances may overestimate the importance of rare sequence variants on community structure (Bálint *et al*. 2016). Different filtrates were included as distinct samples into the analyses since we wanted to see the variation among closely related samples (e.g. the three filtrates taken at the same point of a pond) *versus* the variation among distinct samples (e.g. originating from different ponds). We assumed a negative binomial distribution for both boral and mvabund analyses since read abundances are overdispersed counts, with a strong uniform prior on the dispersion parameter in boral (0-3). The surveyed ponds were markedly different in size, vegetation, water depth, etc, thus we considered pond identity as a good predictor of community composition. The pond effect was evaluated with analysis of variance (likelihood-ratio test, PIT-trap resampling, 1 000 bootstrap replicates). We performed a model-based ordination also for the VAES presence/absence data (since no abundance data was collected during the VAES), and then used a Procrustean superimposition (Peres-Neto & Jackson 2001) to evaluate how the VAES-based ordination of ponds matches the ordination of centroids of eDNA samples. We compared the efficiency of filter preservation (CTAB or dried) on the successful detection of species with a site occupancy model (MacKenzie *et al*. 2002; Bailey *et al*. 2014), implemented in the R package ‘unmarked’ (Fiske *et al*. 2011). We used the single season false-positive occupancy model developed by Miller *et al*. (2011, for details see Site occupancy models in Supplemental information).

For cost comparisons, we considered a typical eDNA survey scenario: samples are collected in the field and later processed in a dedicated laboratory. For the encounter-based survey we considered a scenario with a similar separation of the fieldwork and species identification: species records (audio or visual) are collected by a field biologist, and later identified or verified by an expert in the office/laboratory. The parameters in our cost models are an approximation of the variables involved in the present study and involve a learning effect in the efficiency of the taxonomic expert (see Cost comparison in the Supplemental information for details). The sampling and identification costs of VAES are dependent both on the number of sampling sites, and the number of species since each species needs to be recorded. During the eDNA survey samples are collected by a field biologist, analyzed in a lab and sequenced by an external provider. The costs of eDNA sampling and identification depend on the number of sites, but not on the number of species. We kept some cost factors constant: the costs for training the frog taxonomic expert and the VAES observers, the costs for building up the eDNA metabarcoding facilities (clean rooms and equipment to perform DNA manipulations, except sequencing), and the databases necessary for the sequence assignment. We assume that travel costs are the same for the two survey types, and that the time necessary to walk between frog observations and eDNA sampling points is the same. All model parameters and calculations are accessible on FigShare (Bálint *et al*. 2017).

## Results

The sequencing resulted in 12 742 273 read pairs (deposited in ENA as PRJEB22113), of which 9 479 299 were identified as complete 16S amplicons. De-replication of the reads (see Supplemental Information: Sequence processing for more information on the steps) resulted in 631 003 unique sequences variants. Only 22 706 of these variants were represented by at least 10 reads and retained for further processing. The sequence cleanup resulted in 14 442 high quality sequence variants, 13 497 coming from the present experiment. Of these, 4 805 sequence variants were taxonomically assigned by the *ecotag* command with the 16S_custom database, and 8 692 with the 16S_EMBL database. The *ecotag* outputs are accessible on FigShare (https://doi.org/10.6084/m9.figshare.5099842.v5), and can be combined and simplified with the R script provided on GitHub (https://github.com/MikiBalint/amphibian-eDNA). The assigned sequence variants represented 8 011 631 sequences. After bionformatic processing with OBITools 561 of these sequence variants were identified as ‘head’ sequences, i.e. sequences that have no variants more abundant than a predefined percentage of their own count, here 5% (see also Sequence processing in the Supplemental information) in at least one sample (6 965 866 reads). The original read numbers were re-assigned to these head sequence variants, and the read numbers were used in downstream analyses. Several of these 561 head sequence variants were found also in the negative controls. After removing the reads assigned to these, the final sample - sequence variant abundance matrix contained 5 815 014 sequences. These belonged to several groups: frogs (2 158 534), fish (1 692 613), insects (304 059), mammals (14 006), birds (967), and bacteria and other groups (1 063 156).

Read numbers assigned to frog species varied considerably among the spatial replicates of the eDNA samples (see species_abundance_matrix.csv – zipped - on FigShare: https://figshare.com/articles/_/5099842). Numbers of frog species detected by both methods, eDNA and VAES, varied between 3 and 18 per pond (Table1). Several of the species present at the ponds had no aquatic life phase during the time of survey (i.e. no larvae or adults in the water), and this likely resulted in lack of detection. To better evaluate the performance of the eDNA approach we thus assembled “reduced” VAES data sets containing only those species that are known to have an aquatic life phase during the survey period. For example, we excluded *Leptodactylus fuscus*, a species that was observed calling from the shore of the pond during the survey, but for which no tadpoles were recorded at that time.

**Table 1.** Numbers of detected species by method per pond (VAES = Visual-audio encounter surveys. Reduced VAES = only considering species that were in contact with water during survey).

| Pond | eDNA | VAES | Reduced VAES | Tadpole survey |
| --- | --- | --- | --- | --- |
| T1 | 10 | 8 | 6 | 4 |
| T2 | 7 | 4 | 4 | 3 |
| T3 | 3 | 7 | 3 | 0 |
| T4 | 9 | 15 | 13 | 1 |
| T5 | 13 | 18 | 14 | 1 |

In summary, in the five ponds 31 frog species were detected in total with both VAES and eDNA. Each of these methods detected 25 species, while the TS detected 4 species; 19 species were detected by both eDNA survey and VAES (Fig. 2A; Table 1). Six species were detected only by eDNA, and six species were detected only by VAES. If we consider only species with aquatic life phase during the sampling (“reduced” VAES data set; N=20 for all ponds), of these eDNA detected 19 (Fig. 2B, Table 1, Supplemental Table S1).

**Fig. 2.**
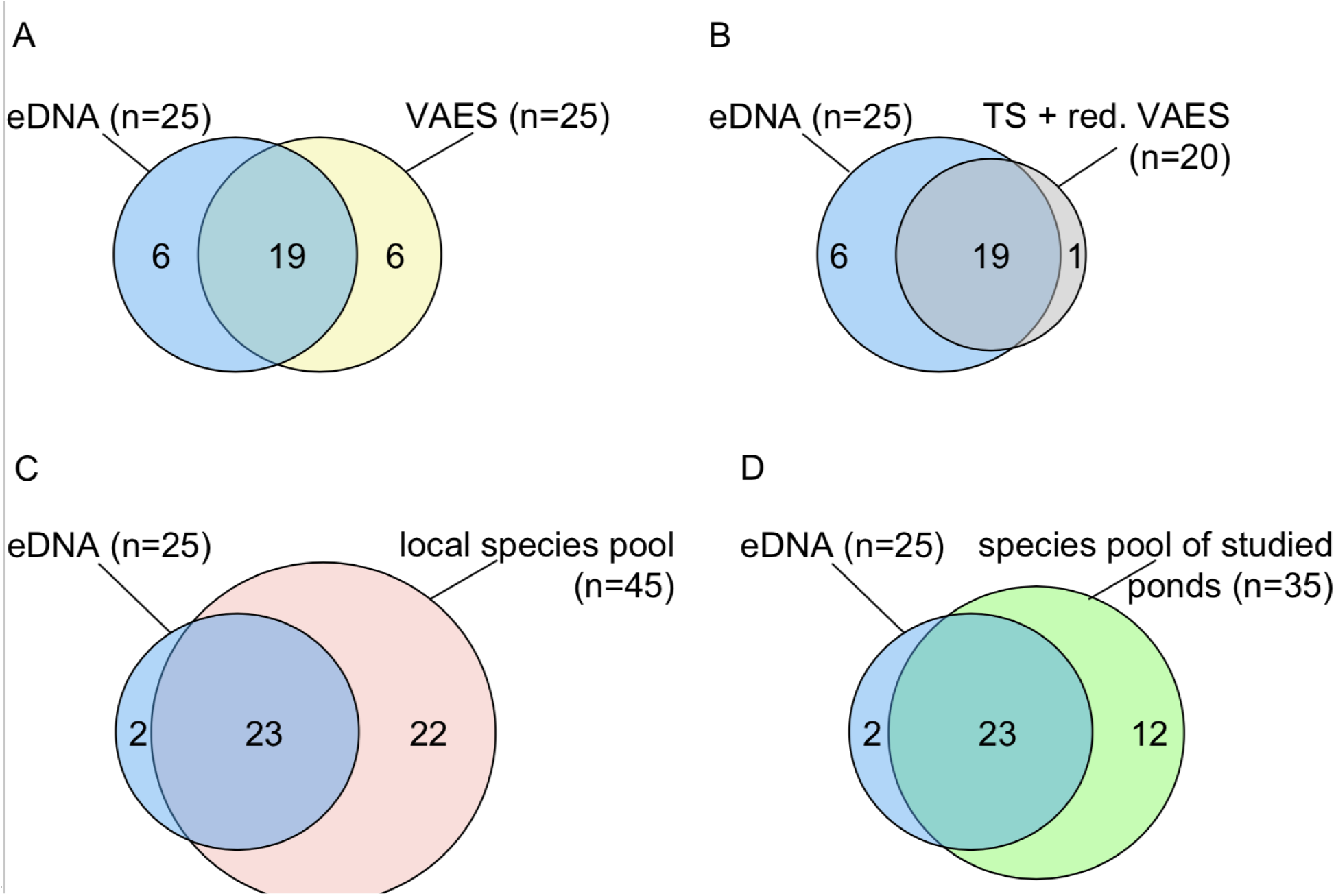
Comparison of species lists generated by eDNA (blue) and A) visual and audio encounter surveys (VAES, yellow), B) tadpole survey (TS) plus reduced VAES - only species with high eDNA detectability in ponds (see text for details; grey), C) complete local species pool (red), and D) species pool of studied ponds based on long-term monitoring (green).

We detected 11 species (of the twelve) from the equimolar DNA concentration positive controls, and 6 in the non-equimolar DNA concentration controls (Supplemental Fig. S2). Read numbers in the equimolar PCR controls were highly variable, but strongly correlated among the controls (*R* > 0.7 for each pair of positive controls, Supplemental Fig. S2A). The read numbers in the non-equimolar positive control were strongly linked to the DNA template concentrations of the PCRs (Pearson correlation coefficient *R* = 0.97, Supplemental Fig. S2B). Only *Leptodactylus vastus* was recovered in the field positive control (water from an aquarium with *L. vastus* tadpoles).

We found no difference in detection probability between the filter preservation methods (CTAB buffer - A: detection probability p_1_=0.278 [0.242; 0.317]; dry - B: p_1_=0.243 [0.209; 0.281]), Supplemental Fig. S3). Interestingly, subsequent filtering of water filtered through glass fiber filters with a 0.2 μm nylon filter - C, appeared to show an increased detection probability (p_1_=0.327, 95% C.I. = [0.285; 0.372]. The three approaches were equivalent with respect to false positive probabilities (Supplemental Fig. S3).

Regarding the community structure assessment based on the eDNA data, replicate samples of each pond grouped relatively tightly on the latent variable model ordination (Fig. 3A). The pond identity was a statistically significant predictor of frog communities in the five ponds (ANOVA, df = 6, dev = 534.99, p < 0.01). This is reflected in the 95% confidence intervals of the group centroids on the ordination which clearly separates all ponds except T1 and T3 (Fig. 3A). The ordination of the eDNA pond centroids closely corresponds with the ordination of observations from the five ponds with VAES (Procrustes permutation test, R = 0.8, p = 0.03, Fig. 3B).

**Fig. 3.**
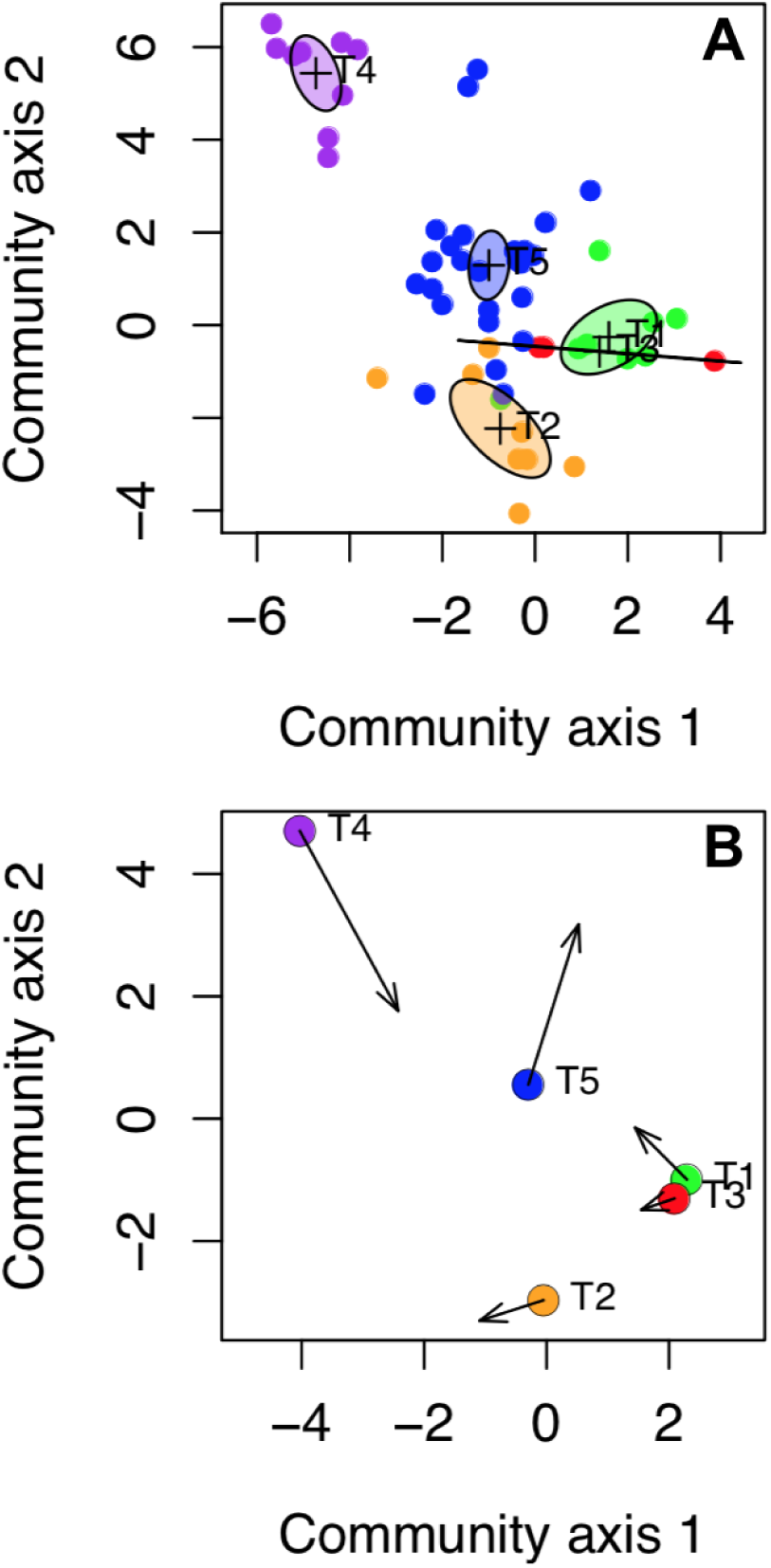
Comparison of ecological signal between visual and audio encounter surveys (VAES) and eDNA metabarcoding surveys. A - Latent variable model ordination of the pond samples according to the read numbers of species. Water samples from each pond were taken at three locations. For ponds T1-T4, three samples at each location were filtered through two GFF (A,B), and one nylon filter (C), resulting in nine samples per pond. For pond T5, each filtration was done three times. Not all samples contained reads after the bioinformatic quality filtering and these samples are not shown on the ordination (see e.g. T3 - red). The ellipses represent 95% confidence intervals for the standard errors of the pond centroids (marked with +). B - Non-metric multidimensional scaling plot of the five ponds (Jaccard distance), according to VAES species presence-absences. The arrows represent the Procrustes rotation of the VAES pond ordination and they target the group centroids of the latent variable model ordination.

The cost model of VAES and eDNA showed that the starting costs (i.e. with few sampling sites) for VAES are relatively low, but these costs rapidly increase until the taxonomic expert becomes familiar with the regional frog fauna (Fig. 4). The VAES price is dependent on the species richness: first, the VAES observer needs to record each species on the field, and then the taxonomic expert needs to listen to each recording (Fig. 4). eDNA metabarcoding has a relatively high entry price since consumables and sequencing are costly, regardless of the number of sites. eDNA survey prices are then a linear function of the number of sampling sites, and an increase in the site numbers simply adds to sampling and consumable costs, but does not influence neither the time spent in the laboratory, nor the sequencing costs (Fig. 4).

**Fig. 4.**
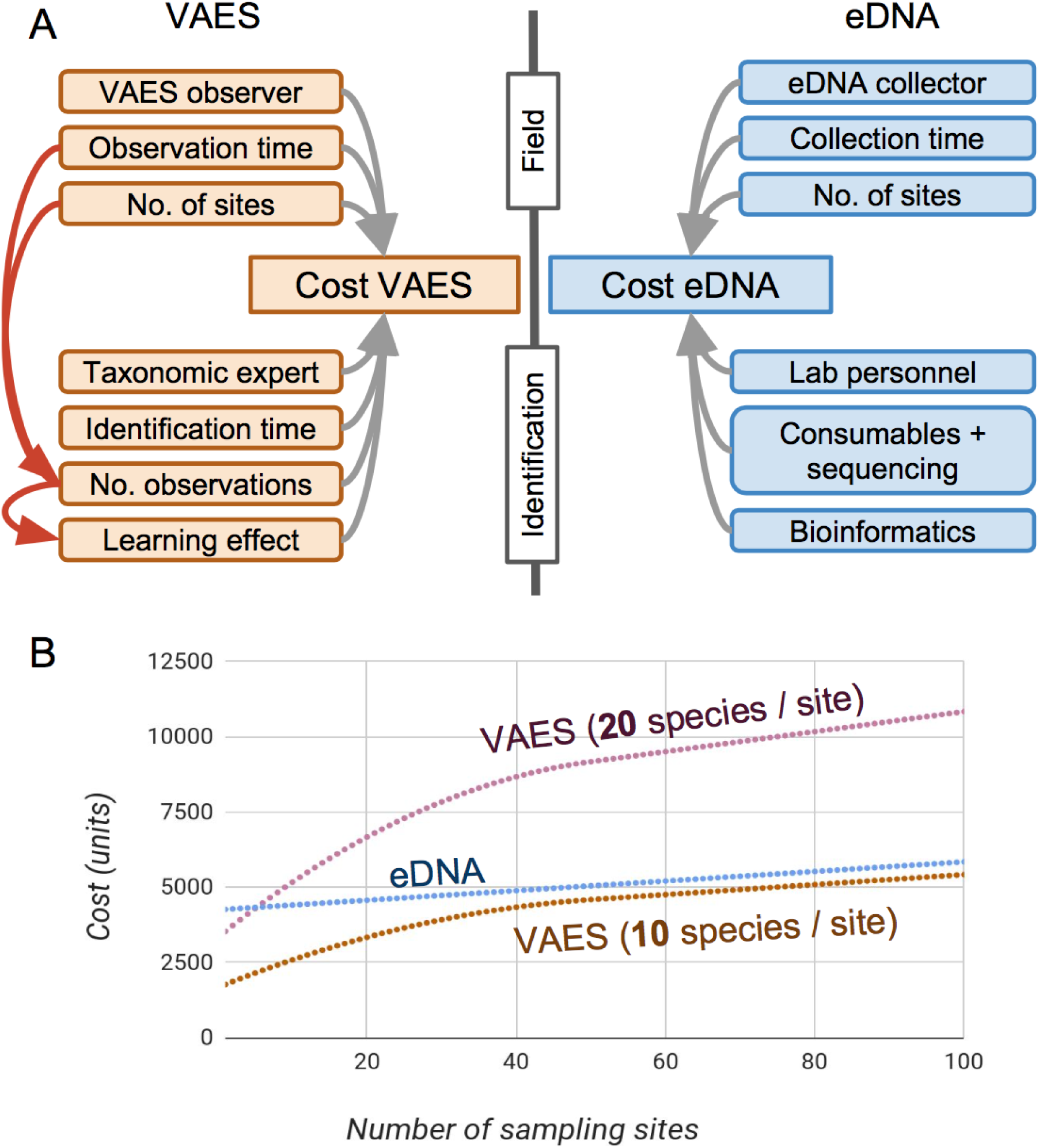
Cost comparisons between visual and audio encounter surveys (VAES) and eDNA metabarcoding surveys. A - Schematic overview of the visual and audio encounter survey (VAES) and eDNA cost models. Model parameters and calculations are accessible through FigShare (Balint_et_al_survey_cost_calculations.xlsx, x). B - Cost comparisons of frog diversity surveys with eDNA, and two VAES scenarios (a low local species richness scenario - 10 species per site, and a medium-high species richness scenario - 20 species per site).

## Discussion

The one-time eDNA survey detected 25 species in the studied ponds and confirmed the presence of (i) almost all species that were in contact with water during the survey time, (ii) all species that had tadpoles in the ponds, (iii) more than half of all 45 species ever recorded from the area, and (iv) about 65% of the 35 species ever recorded in the five surveyed ponds. The eDNA dataset recorded clear differences among the frog communities in the surveyed ponds, a result also in accordance with the VAES data. The comparison of VAES and eDNA cost models show that eDNA biodiversity surveys may be a cost-efficient alternative to VAES in species-rich areas, but not necessarily in areas with low species numbers. Specific recommendations and technical remarks are presented in the Supplemental information.

eDNA approaches for routine use in tropical regions will benefit from the implementation of straightforward and robust sampling techniques. The comparison of the filtration-preservation approaches shows that filters can be dried in the field before being sent to a lab. Currently only few comparative studies exist regarding the preservation of eDNA filtrates on filters (e.g., Hinlo *et al*., 2017; Spens **et al*.*, 2017). Hinlo *et al*. (2017) showed that the simple refrigeration of filters may be preferred to frozen storage. Here we show that filters can be dried in the field: this simplifies transportation and storage since no special precautions are needed, unlike for liquid handling. Interestingly, small pore-size nylon filters seemed to recover a high proportion of target DNA that had previously passed through a 2 μm mesh sized glass fibre filter. While this result may be caused by the different extraction methods used, we consider it likely that a large proportion of DNA occurred in extracellular form. This result suggests that alternative filtering methods based on fine-scale mesh sized filtration with limited filtering volumes may further increase success rates in eDNA studies on amphibian communities. By limited volumes we do not suggest sampling only a few hundred ml of water, but rather to sample enough from a waterbody to ensure representativeness, and then filter from this well-mixed sample until the filter clogs.

During the one-time surveys, six species were only detected with eDNA, and not with VAES (Fig. 2A). Four of these species (*Dermatonotus muelleri, Leptodactylus elenae, Osteocephalus taurinus, Rhinella schneideri*) are quite common in the area. The detection of these four species by eDNA but not by VAES may result from the low abundance of adults or tadpoles in the ponds, or a lack in acoustic activity. In some cases eDNA detections are even likely based on single tadpoles or single adults only. For example, one adult *L. syphax* was heard calling from a crevice of a rocky outcrop, its usual habitat (de Sá *et al*. 2014), approximately 30 m away from T2, but detection by eDNA most probably was due to the presence of tadpoles in pond T2. All other species that had tadpoles in the ponds were also detected by eDNA (Table 1). Furthermore, our results suggest that detection of rare or solitary species by eDNA is possible, since the presence of *L. vastus* in the eDNA most possibly can be traced back to single, territorial adult males that were present during VAES of T4 and T5. These examples show the great potential of eDNA for detection of elusive life stages like tadpoles and less abundant species. The remaining two species detected only with eDNA are not known to occur in the area, thus they are candidates for false assignments. The low read numbers assigned to these species also supports this (Supplemental Table S1). Both of these were assigned with sequence data from EMBL (*L. latinasus*: KM091595, *L. laevis*: AY843696). *Lysapsus laevis* would be the first record of this genus in the study area, but the identification of *Lysapsus* populations only using short 16S eDNA sequences is questionable, especially when considering the unclear taxonomy of the group in Bolivia. Nevertheless, based on their distribution and ecology all of the four known species of *Lysapsus* (*L. boliviana, L. caraya, L. laevis, L. limellum*) may occur at the site (De la Riva *et al*. 2000; Lavilla *et al*. 2004; Reichle 2004a; b; Angulo 2008; Jansen *et al*. 2011; Frost 2016). They may have remained undetected thus far simply because of their small size and rather inconspicuous advertisement call. *Leptodactylus latinasus* is another species possibly occurring in the area that had so far never been recorded (de Sá *et al*. 2014) and we cannot exclude the possibility that the corresponding sequence variant is actually an erroneous variant of one of the six local *Leptodactylus* species.

Species’ ecology might explain why some species were detected only with VAES, but not by eDNA (Fig. 2A, Supplemental Table S1). None of these species are strictly bound to ponds in the life stages occurring during our sampling: *Dendropsophus arndti* and *D. salli*, usually call from plants on the pond shores (Schulze *et al*. 2009) and have only sporadic contact with water. *Leptodactylus fuscus* and *Pseudopaludicola* sp. also do not enter the water but usually call from nearby muddy grounds or grasslands. We did not detect tadpoles of these species in the ponds: the tadpoles of these species develop during the rainy season which triggers reproduction, but our sampling slightly preceded the rainy season. Some of the other undetected species have terrestrial or quasi-terrestrial life histories: *L. fuscus* deposits eggs within foam nests in underground burrows at some distance from ponds and these are washed into nearby water bodies by floods that follow heavy rains (Heyer 1978; Lucas *et al*. 2008). *Boana geographica* was likely missed as a result of PCR bias: we could never recover it from positive controls when DNA from other species was also present (Supplemental Fig. S2), although single-template PCR reactions worked. *H. geographicus* was also not detected by VAES, so it is possible that the species was not present during the survey. If we consider the specific life histories, the one-time eDNA survey only missed a single species (*Sphaenorhynchus lacteus*) in the area which was found calling from the water during the VAES (Fig. 2B, Supplemental Table S1). Regarding the whole local species pool (45 species in 10 years), some of the species (10) may not be detectable by eDNA since they reproduce outside of water (e.g. *Leptodactylus mystacinus*), or are forest dwellers (e.g. *Leptodactylus* cf. *didymus*, Fig. 2D, Table S2). Overall, the results suggests that eDNA performed similarly well in detecting species as VAES, but it was not successful to identify all species occurring at the specific ponds. Differences among individual ponds may result from the differential observational biases of eDNA and VAES approaches, none of which is providing a “true” list of species (hence the long-standing need to model species occupancy). eDNA methods seem to be a powerful tool for the detection of elusive life stages and less abundant species in tropical communities: with a single sampling eDNA detected more than half of the 45 species known to be present in the area, and about 65% of the detectable species from the area (23 out of 35).

The sampling was done at the beginning of the rainy season when only few species reproduced in the ponds. Repeating the sampling during the rainy season could have increased species detections by eDNA. The results show the importance of life history in the metabarcoding-based survey of tropical frogs and emphasize that sampling at multiple time points may be essential for more complete species pools also with eDNA (see Recommendations and technical remarks in the Supplemental information). Comprehensive and taxonomically sound sequence databases are essential for eDNA metabarcoding studies: we had a database that contained all 45 species that were ever recorded in the area. This database was essential for both initial primer development, and taxonomic assignments: indeed, both eDNA-recorded taxa that were not present on the complete local species list of the present study (Fig. 2, Supplemental Table S2) were identified in the EMBL-based assignment that followed the assignment with the custom local 16S database. The importance of sequence assignment databases is long recognized, with considerable efforts underway to establish them (Ratnasingham & Hebert 2007; Coissac *et al*. 2016). There is an urgent need of further DNA sampling to create regional reference data banks, but only a few studies aim at completing DNA sampling of local anurofaunas in South America (e.g. Jansen *et al*. 2011; Guarnizo *et al*. 2015; Moraes *et al*. 2017). However, a preferably complete geographical and taxonomic coverage is necessary for continent-wide eDNA-based frog inventories. Increased DNA sampling will also increase knowledge about actual distributions and taxonomy of Neotropical frogs (including cryptic lineages). This will improve IUCN evaluations, which currently clearly lack distributional information (Supplemental Table S1).

There was considerable variation in the species recorded with the spatial replicates of the eDNA samples, and this underlines that eDNA sampling should be replicated for a good representation of community composition. These samples may be pooled before DNA extraction to optimize costs if site-level variation is not of interest. Interestingly, community structures across ponds inferred from both eDNA and VAES datasets where highly similar (Fig. 3). The eDNA data did not distinguish the communities from ponds T1 and T3, and these communities also grouped closely in the VAES results. Although the comparison of frog community structures among the assessed ponds may be confounded by the different sample sizes (with the eDNA ordination based on many spatial replicates and the VAES ordination on a single observation event per pond), and the eDNA ordinations are based on read-abundances in contrast to the VAES presence-absence ordinations, the correspondence of the results is still striking. Similar results were found in insect (Ji *et al*. 2013; Elbrecht *et al*. 2017) and plant communities (Niemeyer *et al*. 2017) where eDNA-based and morphology-based identifications result in similar conclusions about community structure, despite markedly different species lists. Our results provide further evidence that eDNA-based biodiversity surveys are highly sensitive to differences among ecological communities. These inferences are comparable to those derived with encounter-based observations and are informative about processes that underlay community assembly.

Although central to deciding on a method for biodiversity surveys, cost comparisons are not straightforward since they need to be based on expert knowledge both in VAES and eDNA. Cost comparisons were performed for single species eDNA detection (Huver *et al*. 2015; Davy *et al*. 2015; Smart *et al*. 2016), but we are not aware of frameworks suitable for eDNA metabarcoding. Here the VAES cost estimation is informed by over a decade of field and integrative taxonomic work with tropical frogs (Jansen *et al*. 2007, 2016; Schulze *et al*. 2009; Brusquetti *et al*. 2014), while the eDNA part is only informed by a few years of eDNA biodiversity surveys (Bálint *et al*. 2016; Vörös *et al*. 2017). Such comparisons are urgently needed due to stakeholder demands in eDNA (e.g. governmental agencies, conservation NGOs, fisheries, etc.).

The two species richness scenarios we defined for cost comparisons (Fig. 4) shows that VAES costs become high in regions with high frog richness since they are a function of both the number of sites and the number of species. Biodiversity surveys with eDNA are not necessarily cheaper in low-richness regions since entry costs for the eDNA work are high: lab consumables and sequencing are costly. However, eDNA survey costs are not dependent on the local biodiversity since metabarcoding can consider thousands of species simultaneously in a sample (Taberlet *et al*. 2012b). Consequently, eDNA costs are function of sample numbers, which influence the collection time spent on the field and consumables. eDNA metabarcoding operations are easily scaled up in a sense that hundreds of samples can be simultaneously processed (Ficetola *et al*. 2015).

Several aspects of our cost models are contentious. One issue is whether the relatively untrained VAES observers, or taxonomic experts perform the fieldwork, since taxonomic experts may identify many of the targeted frog species immediately on the field. Currently, most surveys of high-diversity areas are directly done by experts interested in the local fauna, but we argue that this will not work for continental - global biodiversity surveys simply because there are not enough taxonomic experts (Buyck 1999; Haas & Häuser 2003). We also did not consider a scenario when VAES surveys are performed with automated recording devices (ARDs), and sounds are automatically identified by algorithms (see Recommendations and technical remarks in the Supplemental Information). The sound complexity in tropical environments currently prohibits the use of automated sound identifications (Campos-Cerqueira & Aide 2016). It is also difficult to compare fundamental infrastructure and training costs (i.e. the establishment of an eDNA laboratory versus the training of taxonomic experts). Discussions about cost models are timely since they will play major roles in devising much needed continental and global biodiversity surveys (Tittensor *et al*. 2014).

## Conclusions

eDNA seems to be suitable to standardized biodiversity surveys of frogs even in species-rich areas, but it may be overly costly for smaller studies in low-richness regions. Differences between eDNA and traditional surveys seem to result largely from different observational biases. Consideration of life histories promises to improve comprehensiveness of both types of surveys and thus also their correspondence. The eDNA data is certainly suitable to characterize not only community composition, but also the factors that shape communities: this gives an unprecedented opportunity to incorporate eDNA as a standard toolkit for community ecology and macroecology. Assessing community structure with eDNA-based community data foresees global biodiversity surveys and monitoring that will support both biodiversity research, and informed decisions on sustainable use of biological diversity.

## Acknowledgements

The authors thank Jenny Wertheimer for technical assistance during eDNA extraction, José Ribeiro for a helpful discussion on passive acoustic monitoring, and Simon Vitecek for helpfuls suggestions on the manuscript. MJ, AS and JLA thank the owners of Hacienda San Sebastián (family Werding) for their invitation to conduct herpetological surveys on their properties and for logistic support. TC was supported by the USGS Amphibian Research and Monitoring Initiative. MJ was supported by the Erika and Walter Datz-Stiftung, Bad Homburg v. d. H., equipment was sponsored by Globetrotter, Frankfurt. This research presents an outcome of the Centre for Translational Biodiversity Genomics (LOEWE-TBG) and was supported by the research funding programme “LOEWE – Landes-Offensive zur Entwicklung Wissenschaftlich-ökonomischer Exzellenz” of Hesse’s Ministry of Higher Education, Research, and the Arts.

## Data accessibility

DNA sequences: EMBL accession PRJEB22113

Cost model parameters and calculations: FigShare, https://doi.org/10.6084/m9.figshare.5099842.v5

Input data for statistical analyses: FigShare, https://doi.org/10.6084/m9.figshare.5099842.v5.

R-script for bioinformatic filtering and statistical analyses: https://doi.org/10.5281/zenodo.1294092

## Author contributions

M.B., M.J., C.N., C.W designed research; J.L.A., M.B., M.J., O.M., A.S., C.W., B.C. performed research; B.C., T.C., C.N., S.U.P. contributed reagents or analytic tools; M.B., M.J. analysed data; M.B., T.C., M.J., O.M., C.N., S.U.P., wrote the manuscript.

